# Spontaneous control of SIV replication does not prevent immune dysregulation and bacterial dissemination in animals co-infected with *M. tuberculosis*

**DOI:** 10.1101/2021.05.10.443538

**Authors:** Ryan V. Moriarty, Mark A. Rodgers, Amy L. Ellis, Alexis J. Balgeman, Erica C. Larson, Forrest Hopkins, Michael R. Chase, Pauline Maiello, Sarah M. Fortune, Charles A. Scanga, Shelby L. O’Connor

**Affiliations:** Department of Pathology and Laboratory Medicine, University of Wisconsin-Madison, WI 53711; Department of Microbiology and Molecular Genetics, University of Pittsburgh, Pittsburgh, PA 15260; Center for Vaccine Research, University of Pittsburgh, Pittsburgh, PA 15260; Department of Immunology and Infectious Diseases, Harvard T.H. Chan School of Public Health, Boston, MA 02115

## Abstract

Individuals infected with both HIV and *Mycobacterium tuberculosis* (Mtb) are more likely to develop severe Tuberculosis (TB) disease than HIV-naïve individuals. To understand how a chronic pre-existing Simian immunodeficiency virus (SIV) infection impairs the early immune response to Mtb, we used the Mauritian cynomolgus macaque (MCM) model of SIV/Mtb co-infection. We examined the relationship between peripheral viral control and Mtb burden, Mtb dissemination, and immunological function between SIV+ spontaneous controllers, SIV+ non-controllers, and SIV-naïve MCM who were challenged with a barcoded Mtb Erdman strain and necropsied six weeks post infection. Mycobacterial burden was highest in the SIV+ non-controllers in all assessed tissues. In lung granulomas, we found the frequency of CD4+ T cells producing TNFα was reduced in all SIV+ MCM, but CD4+ T cells producing IFNγ were only lower in the SIV+ non-controllers. Further, while all SIV+ MCM had more PD1+ and TIGIT+ T cells in the lung granulomas relative to SIV-naïve MCM, SIV+ controllers exhibited the highest frequency of cells expressing these markers. To measure the effect of SIV infection on within-host bacterial dissemination, we sequenced the molecular barcodes of Mtb present in each tissue and characterized the complexity of the Mtb populations. While Mtb population complexity was not associated with infection group, lymph nodes had increased complexity when compared to lung granulomas across all groups. These results provide evidence SIV+ animals, independent of viral control, exhibit dysregulated immune responses and enhanced dissemination of Mtb, likely contributing to the poor TB disease course across all SIV/Mtb co-infected animals.

**Importance:** HIV and TB remain significant global health issues, despite the availability of treatments. Individuals with HIV, including those who are virally suppressed, are at an increased risk to develop and succumb to severe TB disease when compared to HIV-naïve individuals. Our study aims to understand the relationship between SIV replication, mycobacterial growth, and immunological function in the tissues of co-infected Mauritian cynomolgus macaques during the early phase of Mtb infection. Here we demonstrate that increased viral replication is associated with increased bacterial burden in the tissues and impaired immunologic responses, and that the damage attributed to virus infection is not fully eliminated when animals spontaneously control virus replication.

## Introduction

Human immunodeficiency virus (HIV) and Tuberculosis (TB) remain significant global health burdens, despite the wide availability of treatment. While over 90% of individuals infected with *Mycobacterium tuberculosis* (Mtb) have latent TB infection [1], meaning they do not have symptoms and cannot spread the disease, the co-endemic nature of HIV and TB disease adds another level of complexity. In fact, individuals infected with HIV are still at increased risk of developing and succumbing to active TB disease [2] [3], regardless of whether antiretroviral therapy (ART) has successfully controlled HIV viremia [4–6]. However, the reasons underlying why HIV+ individuals exhibit increased susceptibility to TB disease, independent of their chronic viral load set point, remains a mystery.

The dual susceptibility of nonhuman primates (NHP) to both Simian immunodeficiency virus (SIV), an NHP analog of HIV, and TB has additionally allowed researchers to further probe how the immune system simultaneously combats these pathogens within individual tissues over time. Importantly, the utilization of animal models of co-infection has allowed researchers to control the dose, route, and timing of infection. Often, humans with both HIV and TB are not sure which pathogen they were infected with first or the time between initial infection and co-infection. This uncertainty is related to the high rate of asymptomatic latent TB and the initial non-distinct flu-like symptoms of HIV. While most NHP studies focus on how latent TB, which is the most common form of TB in humans, is reactivated by a newly acquired SIV infection [7], few focus on how a pre-existing SIV infection impacts the host’s ability to contain and clear a new Mtb infection [8, 9].

In order to examine the effect of pre-existing SIV infection on subsequent Mtb infection, we utilized our previously developed Mauritian cynomolgus macaque (MCM) model of SIV and Mtb co-infection. MCMs have limited MHC genetic diversity, such that nearly all of their MHC alleles can be explained by 7 common MHC haplotypes (M1-M7) [10]. Approximately half of MCM containing at least one M1 MHC haplotype spontaneously control viremia following SIVmac239 infection to ≤10^3^ viral copies/mL of plasma, which is considered standard in the SIV field for viral control [11] [12]. We previously found that all M1+ MCM infected with SIV for 6 months followed by Mtb infection for up to 12 weeks developed rapidly progressive TB disease, independent of whether they were spontaneous SIV controllers [8]. In this same study, we observed more rapid granuloma dissemination in SIV+ animals between four and eight weeks post-Mtb infection when compared to SIV-naïve animals [8]. These prior results suggest that the presence or absence of early anti-mycobacterial host immunity at the sites of Mtb replication contributes to whether the host succumbs to rapid TB disease progression.

Here, we tested the hypothesis that Mtb-infected SIV-naïve MCM would exhibit similar Mtb pathogenesis and host immune responses to SIV/Mtb co-infected MCM who spontaneously controlled viral replication (SIV controllers). We also hypothesized that SIV/Mtb co-infected MCM with uncontrolled viral replication would exhibit increased Mtb pathogenesis and a more damaged host immune response when compared to SIV controllers or SIV-naïve animals at 6 weeks post-Mtb infection. By using a molecularly barcoded Mtb, we were able to identify uniquely tagged Mtb bacilli in each tissue and track bacterial dissemination to test the hypothesis that bacterial dissemination would be increased with uncontrolled viral replication. Based on our previous study described above and results observed in Larson et al 2020 [13], we hypothesized that this 6 week timepoint represented a crucial “tipping point” in which the immune response would either prevent severe Mtb disease or allow for rapid mycobacterial replication and increased dissemination. We specifically examined the viral and mycobacterial burden, mycobacterial dissemination, and the immunologic phenotype of CD4+ and CD8+ T cells in the lymph nodes (LN) and lung granulomas. While our observations regarding viral and mycobacterial burden were consistent with our hypothesis, we found the immunologic data was far more nuanced. Cumulatively, this study elaborates on the highly complex system in which increased mycobacterial burden and impaired immune responses is not solely explained by increased SIV replication.

## Results

### Control of plasma SIV replication mirrors that observed in tissues of SIV/Mtb co-infected animals

We infected one cohort of 8 M1+ MCM with SIVmac239 for 6 months, followed by co-infection with a low dose (3-19 CFU) of a molecularly barcoded Mtb Erdman strain for 6 weeks (Figure 1A). A second cohort of 8 SIV-naïve MCM were infected with the same strain and dose of Mtb for 6 weeks [13] [14]. The clinical outcomes of this study are further described in Larson et al [13]. Animals were necropsied 6 weeks after Mtb infection to characterize the pathogen burden and host immune responses in tissues prior to the development of severe TB pathology (e.g. TB pneumonia) [8].

**Figure 1.**
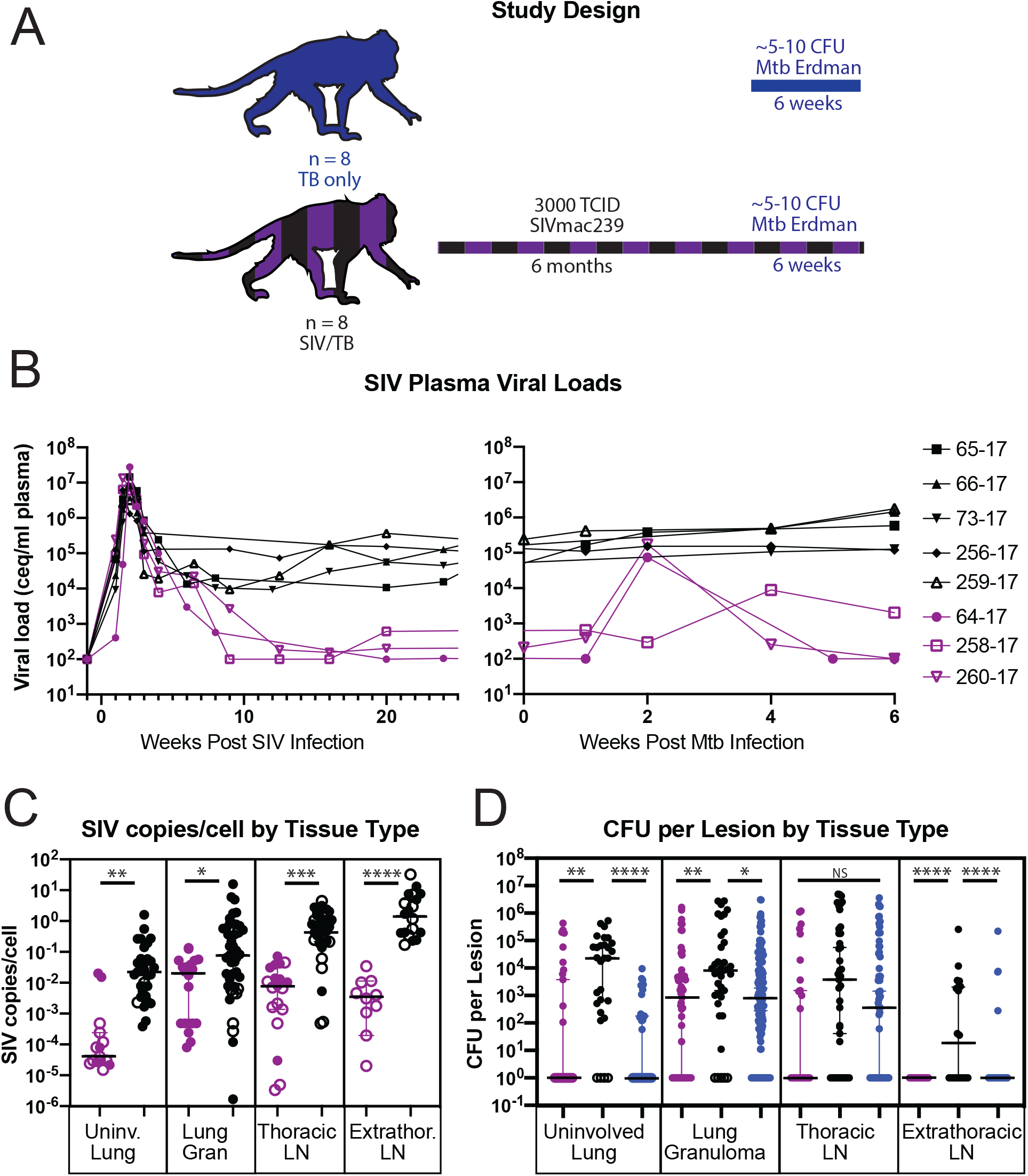
Study design (A), SIV plasma viral loads in our 8 MCM (B), as well as copies/cell (C) and bacterial CFU (D) by tissue type and controller status. Controller status was determined by SIV viral load shown in (A), with purple indicating animals that could spontaneously control SIV replication to less than 1000 SIV copies/mL of plasma, and black indicating animals that could not. Blue dots in figure 1C indicate TB-only animals. Open circles indicate CFU-(sterile) samples. Significance determined by Kruskal-Wallis: *, p < 0.05, **, p <0.01, ***, p < 0.001, ****, p < 0.0001.

SIV plasma viremia was characterized in the SIV-infected cohort (Figure 1B). All animals had at least one copy of the M1 MHC haplotype (Supplemental Table 1), so we expected approximately half the animals would spontaneously control SIV [12]. We found that three SIV+ animals (purple) spontaneously achieved a plasma viral load set point of ≤10^3^ viral copies/mL, and were considered SIV controllers [15] (Figure 1B, purple). The other five animals (black) had a set point of at least 10^4^ copies/mL, which we considered SIV non-controllers. Two weeks after the SIV+ animals were co-infected with Mtb, we detected a transient 2-3 log_10_ spike in viremia in two of the three SIV controllers, but not the SIV non-controllers (Figure 1B). The plasma viremia in these animals spontaneously returned to baseline by 5 weeks post Mtb infection.

Six weeks post Mtb infection, necropsies were performed and individual granulomas, thoracic and extra-thoracic LN, and uninvolved lung tissue were collected and homogenized. Lung tissue was considered “uninvolved” if it did not exhibit macroscopic signs of TB disease at the time of necropsy. However, these samples may harbor Mtb bacilli without overt lesions, so all lung tissue samples were plated for bacterial burden measured by CFU. To assess the viral burden within each sample, SIV *gag* copies/mL were measured in tissue homogenates by qPCR, and SIV copies/cell were calculated as described in the methods. We found that SIV controllers had fewer SIV copies/cell in the majority of their tissue samples compared to SIV non-controllers (Figure 1C). Importantly, because individual tissue samples from a single animal are not truly independent and samples from one animal may overly bias the results, we also compared the median SIV copies/cell per animal between SIV controllers and non-controllers as a more statistically robust measure of viral burden between infection groups. We identified a similar pattern when comparing median viral burden per animal across different tissues, although these differences were not statistically significant (Supplemental Figure 1A).

### Poor control of plasma viremia is associated with increased Mtb bacterial burden in tissues

We compared the bacterial burden in lung tissue, lung granulomas, and LN between SIV controllers, non-controllers, and SIV-naïve MCM. We hypothesized that the SIV controllers (Figure 1D, purple) and SIV-naïve (Figure 1D, blue) MCM would have similar Mtb bacterial burdens, while the SIV non-controllers (Figure 1D, black) would have higher bacterial burden compared to the other cohorts. We found no statistically significant differences in Mtb burden between the SIV controllers and the SIV-naïve MCM for any of the tissues examined (Figure 1D, purple vs. blue). In contrast, the bacterial burden was higher in lung tissue, lung granulomas, and extra-thoracic LN in the SIV non-controllers (black) than in the SIV-naïve (blue) and SIV controller (purple) animals (Figure 1D). Again, we compared the animal median CFU in each tissue type to confirm that a single animal was not overly biasing our lesion-based analysis. While we observed a similar pattern to the lesion-based analysis, the only significant relationship was between the lung granulomas of SIV non-controllers when compared to the SIV-naïve animals. (Supplemental Figure 1B).

We then compared the total Mtb bacterial burden per animal across all three groups. In contrast to our analysis of individual lesions, the total thoracic bacterial burden was no different across groups (Supplemental Figure 2A), similar to what was observed when SIV+ MCM were considered as a single group [13]. We also compared the frequency of sterile (CFU-) lesions between the groups. Regardless of if SIV+ MCM were stratified by controller status, there were no statistically significant differences between the percent of sterile granulomas in SIV-infected MCM when compared to those who were SIV-naïve (Supplemental Figure 2B). This suggests that control of peripheral viremia is not related to the percentage of sterile granulomas present at six weeks following Mtb co-infection.

We then examined if there was a correlation between bacterial burden and viral burden in the same tissues across all SIV+ animals. With the limited numbers of animals and samples, we could not stratify these correlations by tissue type or SIV controller status. As a single group, we found a significant positive correlation between bacterial burden and the amount of virus per cell in individual tissues (Supplemental Figure 3). Taken together with our results from Figure 1, we posit that the presence of actively replicating SIV in a given tissue impairs host control of mycobacterial growth, leading to an overall increase in total tissue CFU.

### Reduced frequency of TNFα-producing CD4+ T in lung granulomas is independent of SIV controller status

The frequency of TNFα- and IFNγ-producing CD4+ and CD8+ T cells from granulomas and lung tissue homogenates were measured as part of our previous study [13, 14]. In that study [13], we found evidence of fewer TNFα-producing CD4+ and CD8+ T cells isolated from lesions of SIV+ animals when compared to SIV-naïve animals. Here, we further stratified the samples into those isolated from SIV controllers and non-controllers.

We first examined the frequency of ex vivo TNFα- or IFNγ-producing CD4+ T cells isolated from lung granulomas and thoracic LN [13, 14]. While there was no difference in the frequency of cytokine-producing CD4+ T cells in the thoracic LN between SIV+ groups, similar to that previously observed by Diedrich et al. in cervical LN biopsies of HIV/Mtb co-infected individuals [16], we found a lower frequency of IFNγ-producing CD4+ T cells isolated from lung granulomas of SIV non-controllers when compared to either SIV controllers or SIV-naïve animals (Figure 2B). In addition, there was also a lower frequency of TNFα-producing CD4+ T cells isolated from lung granulomas of SIV non-controllers when compared to SIV-naïve animals (Figure 2A, B). This suggests that the presence of fewer TNFα-producing CD4+ T cells in lung granulomas of SIV+ animals is independent of SIV controller status, but the lower frequency of IFNγ-producing CD4+ T cells is particularly severe in animals unable to control SIV.

**Figure 2.**
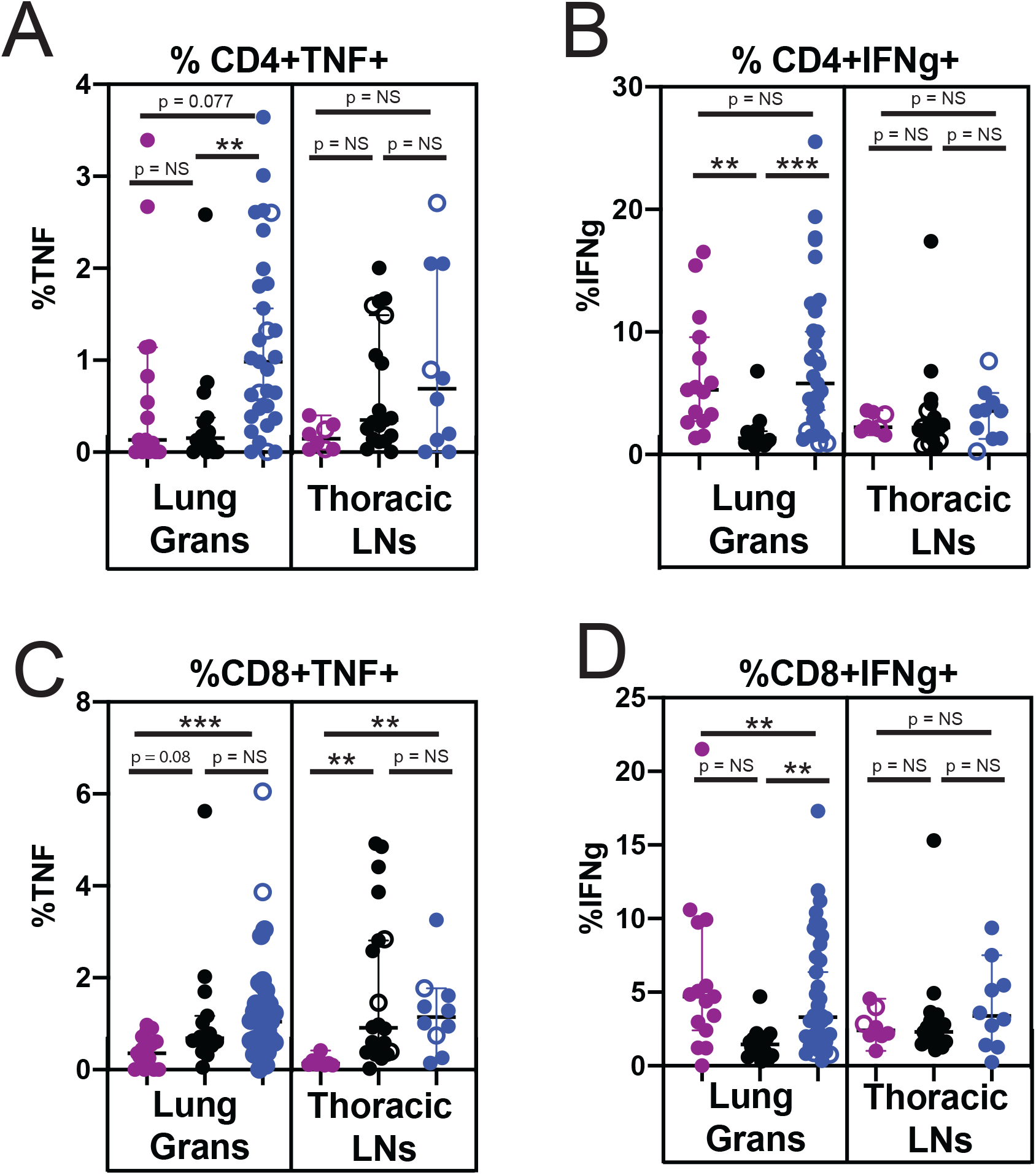
Freuency of CD4 (A, B) and CD8 (C, D) T cells producing TNFa (A, C) or IFNg (B, D) in lung granulomas (left panel) and thoracic lymph nodes (right panel) of spontaneous SIV controllers (purple), SIV non-controllers (black), and TB only (blue) animals. Open circles indicate sterile (CFU-) samples. Significance determined by Kruskal-Wallis: *, p <0.05, **, p < 0.01, ***, p <0.001, ****, p < 0.0001.

*Ex vivo* cytokine-producing CD8+ T cells from the same lesions of all three groups were compared, and the results were more nuanced (Figure 2C, D). The frequency of cytokine-producing CD8+ T cells was lower in the lesions of SIV controllers when compared to those present in the SIV-naïve cohort. However, SIV controllers and non-controllers both had a lower frequency of CD8+ T cells producing IFNγ than SIV-naïve animals in the lung granulomas. In contrast, the frequency of TNFα-producing CD8+ T cells was similar between the SIV non-controllers and SIV-naïve animals. These assays were performed without additional peptide stimulation, so we cannot determine if the CD8+ T cells were specific for SIV or Mtb. However, these data indicate that independent of SIV controller status, the ongoing SIV infection perturbs the cytokine milieu in the lung microenvironment when compared to SIV-naive animals.

As above, these differences are not evident when the median frequency of cytokine-producing T cells per animal is compared across all three groups (Supplemental Figure 4), likely as a result of low animal numbers. Unfortunately, we were only able to characterize the T cell frequencies in two SIV controllers. Further studies and experiments will be required to fully elucidate the relationships between the frequency of cytokine producing cells and systemic control of SIV.

### High frequencies of T cells expressing PD1 and TIGIT were present in the lung granulomas of SIV+ animals, independent of SIV controller status

We previously assessed the frequency of PD1- and TIGIT-expressing CD4+ and CD8+ T cells in thoracic LN and lung granulomas of animals in this study [13, 14]. Similar to the cytokine data, we now stratify these samples by SIV controller status. We found that the SIV-naïve MCM had significantly fewer CD4+ and CD8+ T cells expressing PD1 or TIGIT in both the lung granulomas and thoracic LN when compared to SIV non-controllers, and fewer CD4+ and CD8+ T cells expressing PD1 or TIGIT in the lung granulomas, but not thoracic LN, of SIV controllers (Figure 3). The LN results were not entirely surprising, as HIV replication, and by extension, SIV replication, occurs largely in the LN [17, 18]. PD1+ CD4+ T cells in the LN have been reported to be the primary source of replication-competent HIV [18] and correlate with viral load [19] so it was not entirely surprising that the frequency of T cells expressing PD1 was highest in the SIV non-controller cohort.

**Figure 3.**
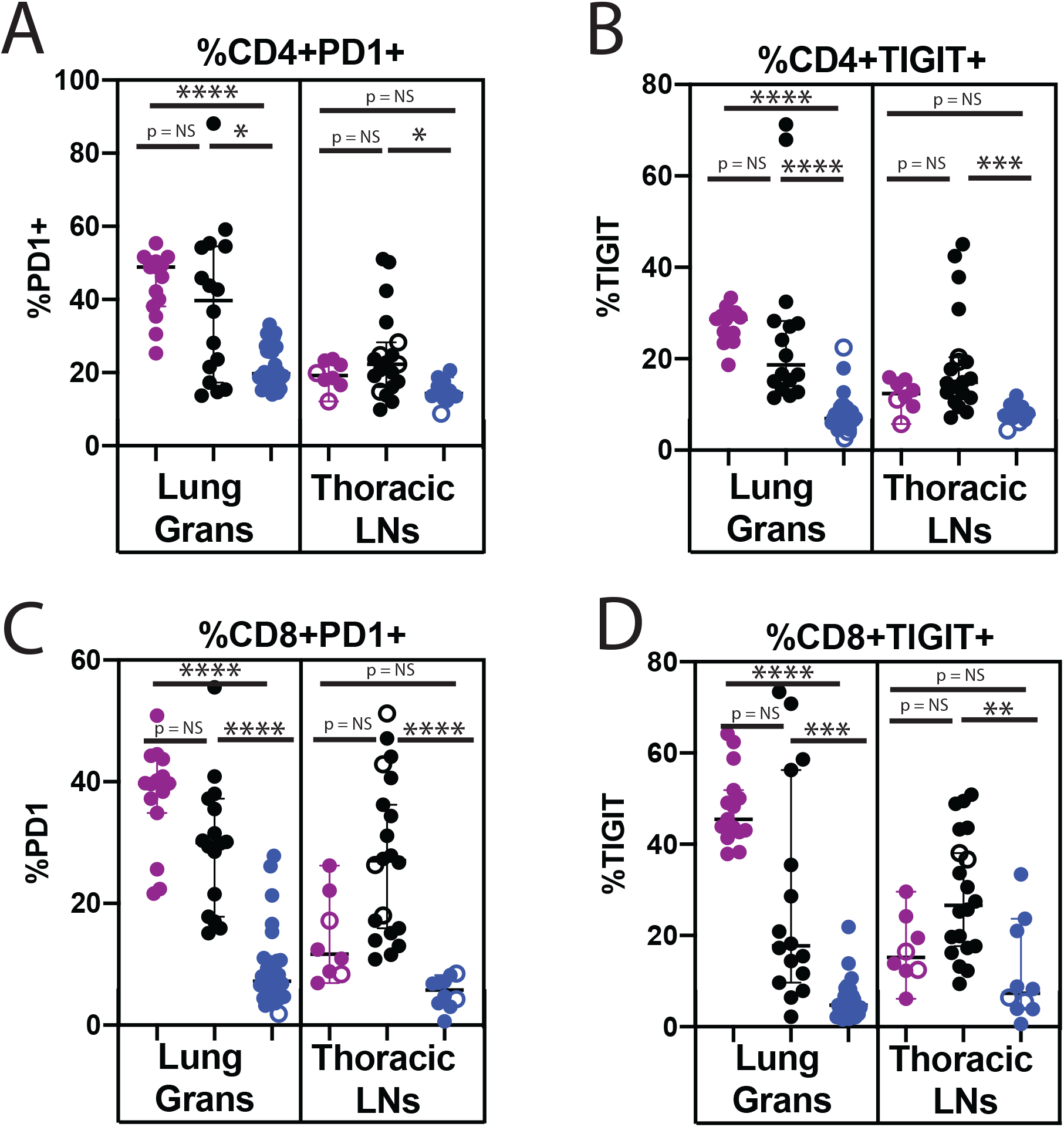
Frequency of CD4 (A, B) and CD8 (C, D) T cells expressing activation markers PD1 (A, C) and TIGIT (B, D) in the lung granulomas (left panel) and thoracic LN (right panel). Spontaneous SIV controllers are shown in purple, non-controllers are shown in black, and TB only animals are shown in blue. Open circles indicate CFU-samples. Significance is determined by Kruskal-Wallis: *, p < 0.05, **, p < 0.01, ***, p < 0.001, ****, p < 0.0001.

Again, we compared the median frequencies of the PD1+ and TIGIT+ T cell populations in each animal, calculated by combining the data from each lesion per animal (Supplemental Figure 5). While there were fewer significant relationships, we identified similar trends to the lesion-based analysis. Even with such a small number of animals, we identified significantly fewer median TIGIT-expressing CD4+ T cells in the thoracic LN of SIV-naïve animals when compared to SIV non-controllers (Figure S5B). Additionally, the SIV-naïve MCM had significantly fewer median PD1-expressing CD8+ T cells in both the lung granulomas and thoracic LN when compared to the SIV non-controllers (Figure S5C).

### Lymph nodes are sites of increased Mtb dissemination

Many prior reports describe increased extra-thoracic bacterial disease in individuals with severe HIV or SIV disease [9] [20] [21]. Therefore, we hypothesized that actively replicating virus in a persistent HIV/SIV infection impairs the host’s ability to control bacterial growth and limit dissemination. We infected animals with a molecularly barcoded Mtb Erdman strain [22], which allowed us to identify the progeny of the uniquely tagged bacterium in each uninvolved lung, LN, and granuloma sample across cohorts at the time of necropsy. This system was previously used in Chinese cynomolgus macaques to show that, in the absence of SIV, granulomas are typically seeded by a single bacterium and these lesions contribute to infection of the draining LN [22, 23]. We used Simpson’s Diversity Index (SDI) and the number and frequency of unique barcoded Mtb in each sample to quantitatively examine the diversity and richness of the bacterial population present. With this metric, tissue samples containing more diverse populations of barcoded Mtb strains would have a value closer to 1, whereas samples with a single barcoded Mtb strain would have a value of 0. This allowed us to quantitatively assess the impact of chronic SIV infection on Mtb dissemination in the SIV/Mtb co-infection model.

Strikingly, we did not identify significant differences in the SDI of the Mtb population in individual tissues as a function of SIV control (Figure 4A), but found that SIV-naïve MCM had a lower SDI of the Mtb population in all tissues combined (indicating more lesions with a single barcoded Mtb strain) when compared to SIV+ MCM. When tissues were examined independently and the SIV+ animals were combined into one group, the SDI of the Mtb population of the SIV+ MCM was significantly higher when compared to the SDI of the Mtb population of the SIV-naïve MCM, except for the thoracic LNs (Figure 4B). Indeed, irrespective of SIV status, the SDI of the Mtb population found in the LN was significantly higher than in the lung granulomas (Figure 4C). Notably, the SDI of the Mtb population isolated from apparently uninvolved lung was only higher than in granulomas in the SIV+ non-controllers (Figure 4C). This likely reflects greater bacterial dissemination in SIV non-controllers, since Mtb was more commonly found in tissues that had otherwise appeared uninvolved at the time of necropsy. These data support the hypothesis that SIV infection, regardless of the level of peripheral viremia, contributes to mycobacterial dissemination, particularly in the lung granulomas and surrounding lung tissue.

**Figure 4.**
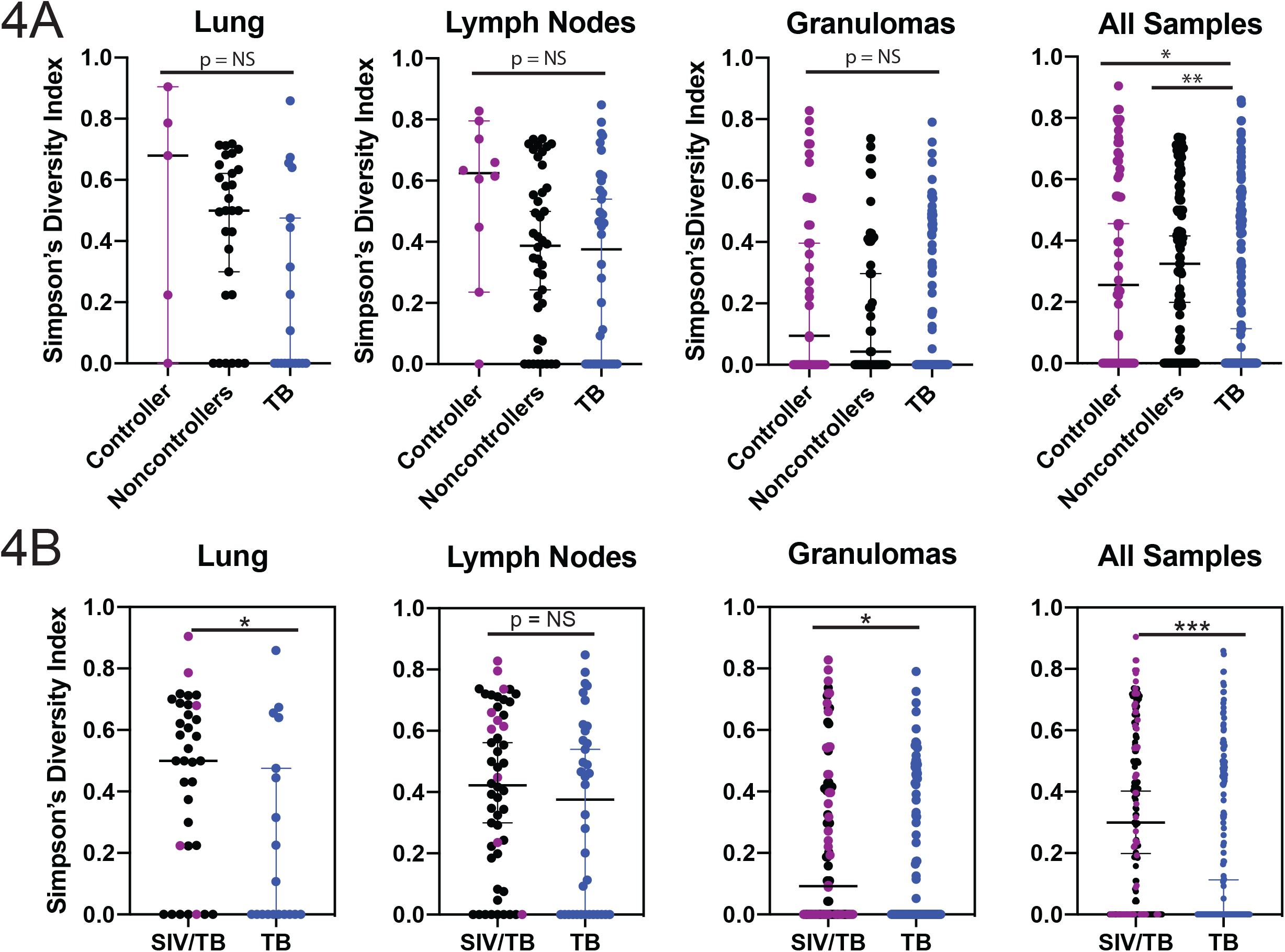
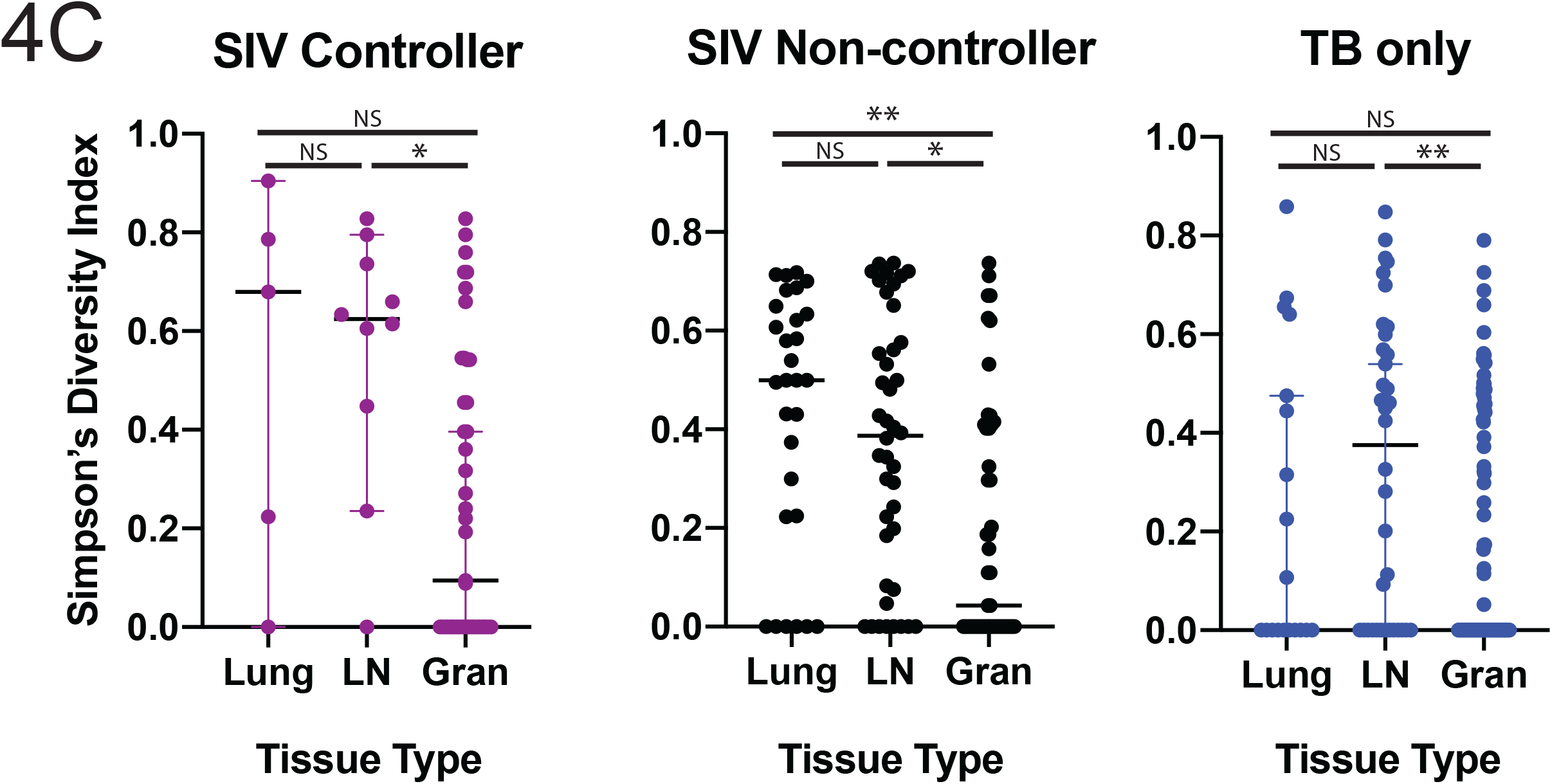
Mtb barcode diversity as calculated by Simpson’s Diversity Index in uninvolved lung, LN, granulomas, and all tissues combined, separated by infection and controller status (A), as well as between SIV+ MCM and SIV-naïve MCM (B). Mtb barcode diversity within each cohort (C). SIV controllers are indicated in purple, SIV non-controllers in black, and SIV-naïve in blue. Significance is determined by Kruskal-Wallis: *, p < 0.05, **, p < 0.01, ***, p < 0.001, ****, p < 0.0001.

## Discussion

In this study, we examined the burden of SIV and Mtb, as well as corresponding CD4+ and CD8+ T cell phenotypes, in the tissues of SIV+ or SIV-naïve MCM infected with Mtb for 6 weeks. This time point was specifically chosen because it was before the pathology was driven by the host immune responses and when we observed more rapid granuloma dissemination between SIV+ and SIV-naïve MCM [8]. The animals and overall host immune responses were described previously [13] [14], but this current study is particularly unique because we could dissect distinct immunological phenotypes between SIV-controllers and SIV-non-controllers. We also provide a detailed analysis of how SIV and Mtb may affect the growth of each other.

Despite control of plasma viremia, we found that the SIV controllers were still susceptible to TB disease, much like HIV+ individuals on ART who remain more susceptible to TB than HIV-naïve individuals [6]. There was a lower frequency of CD4+ T cells in SIV-infected MCM when compared to SIV-naïve animals [13], and this was exacerbated when comparing the SIV non-controllers to the SIV controllers (data not shown). However, while lower CD4+ T cell counts are associated with increased TB incidence, HIV+/SIV+ individuals with high CD4 counts also develop severe TB disease [24]. Our current study is unique because we stratified groups by SIV control to dissect out how this feature influenced whether there were immunological and bacterial differences relating to TB disease course.

We found that co-infection of SIV+ animals with Mtb transiently induced virus replication. Eight MCM were infected with SIV, and three spontaneously controlled viral replication in the plasma without antiretroviral treatment. Among SIV controllers, we observed a transient spike in plasma viremia following Mtb co-infection (Figure 1B). We hypothesize that direct and indirect mechanisms were responsible for this transient spike. CD4+ T cells are vital in the response to Mtb infection, particularly in the lungs [25], and are primarily targets of HIV/SIV infection [21, 26]. Thus, reactivation of latent virus from these resting T cells in the lung could contribute to this observed spike in plasma viremia [20]. Indirect activation of T cells harboring SIV may contribute to this spike in viremia, as the production of TNFα associated with Mtb co-infection can contribute to increased viral replication [27] [28] [29].

We and others have reported increased bacterial burden in the individual lung tissue, lung granulomas, thoracic LN, and extrapulmonary tissues of SIV/Mtb co-infected animals relative to those who were SIV-naïve [8] [9], which is consistent with our findings here. We hypothesized that high SIV viral loads would correlate to increased bacterial CFU in Mtb-affected tissues, and identified a small but highly significant positive correlation between Mtb CFU and SIV viral load across all the SIV+ animals (Supplemental Figure 3). We were not able to determine if the SIV infection was driving Mtb growth, or if Mtb was driving SIV replication in these tissues. However, this does provide evidence that the simultaneous presence of these pathogens leads to exacerbation of both.

As a result of increased viral replication in tissues, we hypothesized that the CD4+ and CD8+ T cell phenotypes in SIV controllers would be more similar to SIV-naïve animals, compared to the SIV non-controllers. Similar to our previous study [13], we found a lower frequency of CD4+ T cells produced TNFα in the lung granulomas, independent of SIV controller status. When we further examined the CD8+ T cells, the SIV controllers had the lowest frequency of CD8+ T cells producing TNFα. These results suggest that SIV-infection, independent of controller status, may be associated with poor TNFα production in the Mtb-affected tissues. TNFα is known to be essential for containment of Mtb replication [30], and it is known that anti-TNFα antibody treatment leads to reactivation of TB disease [28, 31]. Thus, poor TB disease course in all SIV/HIV infected individuals may be, in part, related to the reduced production of TNFα. However, additional studies with SIV+ animals treated with antiretrovirals will be needed to determine whether this routine treatment restores TNFα production at these sites.

We also examined the frequency of IFNγ-producing CD4+ and CD8+ T cells. We found that the SIV non-controllers had fewer CD4+ and CD8+ T cells producing IFNγ in the lung granulomas, when compared to the SIV-naïve animals. Notably, the frequency of IFNγ-producing T cells was less affected in the SIV controllers, when compared to SIV-naïve animals. Unfortunately, we do not know if production of IFNγ was derived from Mtb-specific or SIV-specific T cells, and whether that would affect Mtb containment. IFNγ is an essential cytokine required for Mtb containment [32] and it is also produced by SIV-specific CD8 T cells [33]. How the antigen-specificity of the T cells producing IFNγ affects TB disease in SIV/Mtb co-infected animals is not known. Nonetheless, dysregulated production of IFNγ in SIV+ animals may have contributed to further exacerbation of TB disease in these animals.

Previous reports have indicated that HIV/SIV infected individuals have a higher frequency of cells expressing PD1 and TIGIT [19] [34] [35]. However, our data did not show that the level of peripheral viral replication was related to expression of these markers. We observed no significant differences between the frequency of PD1 or TIGIT-expressing CD4 or CD8 T cells when comparing SIV controllers and SIV non-controllers. Further, both ART-treated and ART-naïve HIV+ individuals often have a higher frequency of PD1-expressing cells compared to their HIV-negative counterparts [36] [37]. While this is consistent with our data in that the SIV-naïve MCM have significantly lower frequencies of PD1-expressing T cells in both the lung granulomas and thoracic LN when compared to SIV+ non-controllers, the thoracic LN of SIV+ controllers have similar frequencies of PD1-expressing T cells to the SIV-naïve MCM. Therefore, the dynamic differences in the frequency of PD1-expressing T cells across tissue types is not solely mediated by the amount of SIV replication in the individual tissue, and additional parameters, such as cytokine production, must be considered.

Individuals with a pre-existing SIV/HIV infection also have a higher risk for developing extrapulmonary TB [9] [20] [21]. To measure bacterial dissemination within a host, we examined the diversity of Mtb barcodes present in uninvolved lung, LN, and granuloma samples from the three cohorts. Interestingly, we did not observe a statistically significant difference between the SDI of the barcoded Mtb population in the lung, LN, or granulomas as a function of SIV control. However, we found that SIV+ MCM had an increased SDI of the barcoded Mtb population in all tissues, besides the LN, when compared to SIV-naïve animals (Figure 4). This may be attributed to the high frequency of extrapulmonary TB that is common in MCMs [38], even when animals remain SIV-naïve. While this is the first study, to our knowledge, to specifically report the effects of viral replication on mycobacterial dissemination in a macaque model, further studies will be required to fully elucidate the mechanisms influencing these results, particularly regarding the differences in barcode diversity in different tissue types.

Cumulatively, this study highlights the importance of understanding the interaction between SIV and Mtb in tissues and the impact of increased viral replication on early TB disease. Because SIV controllers had limited peripheral viral replication, we hypothesize that spontaneous SIV controllers and ART-treated SIV+ MCM have similar immune responses to Mtb co-infection. Future studies will need to directly compare ART-treated and ART-naive SIV+ MCM and examine SIV-specific and Mtb-specific T cell responses in tissues using ex vivo functional assays. Continuing to include barcoded pathogens (both SIV and Mtb) in animal studies, will help improve our understanding of the impact of ART on SIV and Mtb dissemination. Ultimately, the study reported here provides further evidence that there are immunological and microbiological factors beyond control of SIV replication that contribute to early development of TB disease in co-infected individuals. Future studies will also need to evaluate the contribution of viral control on chronic TB disease. Thus, this study will continue to prompt new questions about tissue-specific responses in SIV+ or HIV+ individuals who are co-infected with Mtb.

## Materials and Methods

### Animal care and Ethics statement

Mauritian cynomolgus macaques (*Macaca fascicularis*; MCM) were obtained from Bioculture, Ltd. (Mauritius). All MCM chosen for this study were selected by genotyping to have at least one copy of the M1 MHC haplotype [39] (Supplemental Table 1). Animals (n = 16) were housed in a BSL2+ facility at the University of Pittsburgh (U. Pitt.) during SIV infection and transferred to a BSL3+ facility within the Regional Biocontainment Laboratory for Mtb infection. The U. Pitt. Institutional Animal Care and Use Committee (IACUC) approved all animal procedures and protocols and adheres to guidelines established in the Weatherall report (8th Edition) and the Guide for the Care and Use of Laboratory Animals. The studies described here were conducted under IACUC study protocols 18032418 and 15035401, which were reviewed and approved by the U. Pitt. U. Pitt follows national guidelines established in the Animal Welfare Act (7 U.S.C. Sections 2131-2159) and Guide for the Care and Use of Laboratory Animals (8th Edition) as mandated by the U.S. Public Health Service Polity. U. Pitt’s Animal Welfare Act Assurance Number is A3187-01. Macaques were pair-housed at U. Pitt in caging measuring 4.3 square feet per animal and spaced to allow visual and tactile contact with neighboring conspecifics in rooms with autonomously controlled temperature, humidity, and lighting. Animals had *ad libitem* access to water. Macaques were fed biscuits formulated for nonhuman primates (NHP) twice daily, with additional fresh fruits, vegetables, or foraging mix provided at least 4 days/week. Our NHP enrichment specialist designed and oversaw a three-component enhanced enrichment plan in which species-specific behaviors are encouraged. Animals are provided with access to toys and manipulanda filled with food treats such as frozen fruit and peanut butter, which are rotated regularly. Foraging behaviors are stimulated with puzzle feeders, cardboard tubes containing small food items, and foraging boards placed in the cage. Interactions between cages are stimulated using adjustable mirrors accessible to the animals. Human and macaque interactions are encouraged and occur daily, adhering to established safety protocols. While performing tasks in the housing area, animal caretakers are encouraged to interact with the animals through talking and facial expressions. A strict schedule is followed for routine procedures such as feeding and cage cleaning to allow the animals to acclimate to a daily schedule. Auditory and visual stimulation is provided to all macaques through radios or TV/video equipment showing cartoons or other formats for children in the housing areas. Enrichment is rotated routinely so animals are not repetitively exposed to the same videos or radios. Appetite, attitude, activity level, hydration status, etc. were checked at least twice daily. Animals were monitored closely for evidence of disease (anorexia, lethargy, coughing, dyspnea, tachypnea, etc) following SIV and/or Mtb infection. Physical exams, including weights, were regularly performed. Ketamine was used to sedate animals prior to all veterinary procedures (blood draws, etc). Disease progression was monitored following Mtb infection by monthly PET/CT imaging [40]. Animals were closely monitored for any signs of pain or distress by experienced veterinary technicians, and appropriate medication or supportive care was provided if necessary. Any animal considered to have advanced disease or intractable pain or distress from any cause was sedated with ketamine and then humanely euthanized using sodium pentobarbital (65 mg/kg, IV). A trained veterinary professional confirms death by lack of heartbeat and pupillary responses.

### SIV and Mtb infection of MCM

SIV/Mtb co-infected animals(n=8) were infected intrarectally with 3,000 TCID_50_ SIVmac239. Six months following SIV infection, animals were co-infected with a low dose (3-19 CFU) of a molecularly barcoded Mtb (Erdman strain) [22] via bronchoscopic instillation, as described previously [8]. Animals in the TB only control group (n=8) were infected with barcoded Mtb in an identical manner. Clinical testing and PET/CT imaging was used to monitor TB progression [8]. Animals were humanely euthanized six weeks following Mtb infection. Necropsies were performed using PET/CT images to map individual granulomas and guide their excision. Random lung lobe samples, peripheral LN, and thoracic LN were also harvested [38]; further described in Larson et al., (manuscript in prep).

### Sample collection

Tissues collected at necropsy were homogenized using Medimachines (BD Biosciences) and an aliquot of each homogenate was plated on 7H11 agar plates to quantify Mtb colony forming units (CFU) as previously described [8, 38]. Another aliquot was used for flow cytometry, with viable cells quantified by trypan blue exclusion in a hemocytometer, as described in Larson et al (in prep).

### Flow Cytometry

Flow cytometry was conducted as described previously [14]. Briefly, 1×10^6^ cells (or all cells obtained from granulomas with <1×10^6^ cells) were stained with 0.25 ug of Mamu MR1 5-OP-RU or Ac-6-FP tetramer for one hour in the presence of 500 nM Dasatinib (Thermo Fisher Scientific; Cat No. NC0897653). When TCRVα7.2 co-staining was performed, the antibody was added 30 minutes after the addition of the MR1 tetramer. Cells were washed once with FACS buffer (10% Fetal Bovine Serum (FBS) in a 1X PBS solution) supplemented with 500 nM Dasatinib, then surface antibody staining was performed for 20 minutes in FACS buffer + 500 nM Dasatinib. Antibodies used for surface staining are listed in Table 1. Samples were fixed in 1% paraformaldehyde for a minimum of 20 minutes. For intracellular staining, cells were washed twice with FACS buffer and staining with antibodies was performed in Medium B permeabilization buffer (Thermo Fisher Scientific, Cat. No. GAS002S-100) for 20 minutes at room temperature. For intranuclear staining, the True Nuclear Transcription Factor Buffer Set (Biolegend; San Diego, CA) was used according to manufacturer’s instructions. Briefly, cells were fixed in the TrueNuclear™ fixation solution for 1 hour, then washed three times with the permeabilization buffer. Cells were then stained with the transcription factors indicated in Table 1 at 4°C for one hour, rinsed three times with permeabilization buffer, then resuspended in FACS buffer. Flow cytometry was performed on a BD LSR II (Becton Dickinson; Franklin Lakes, NJ), and the data were analyzed using FlowJo software for Macintosh (version 9.9.3 or version 10.1).

**Table 1.** Antibodies used in staining panels for flow cytometry.

| Marker | Clone(s) | Fluorochrome(s) | Purpose of Use | Surface/Intracellular/intranuclear |
| --- | --- | --- | --- | --- |
| CD45 | D058-1283 | BV786 | Lineage | Surface |
| CD3 | SP34-2 | AF700, BV650 | T cell marker | Surface |
| CD4 | OKT4, L200 | PE Cy7, BV510, BV711, BV786 | T cell marker | Surface |
| CD8 | RPAT8, SK1 | PacBlue, BV510, BV605, BV711, BV786 | T cell marker | Surface |
| CD206 | 19.2 | PE Cy5 | Macrophage exclusion | Surface |
| TCRVα7.2 | 3C10 | BV421, BV605 | MAIT cell marker | Surface |
| CCR5 | J418F1 | BV421 | Chemokine/MAIT lung homing | Surface |
| CCR6 | 11A9 | PE CF594, PE Cy7 | Chemokine/gut homing | Surface |
| CD127 | MB15-18C9 | PE | IL-7 receptor | Surface |
| CXCR3 | G025H7 | PE Dazzle 594 | Chemokine/MAIT lung homing | Surface |
| CCR7 | FAB197F | FITC | Chemokine/lymph homing | Surface |
| CD28 | CD28.2 | APC, BV510 | T cell memory | Surface |
| CD95 | DX2 | PE Cy5 | T cell memory | Surface |
| T-Bet | 4B10 | BV605 | Transcriptional marker | Intranuclear |
| Eomes | WD1928 | PE Cy7 | Transcriptional marker | Intranuclear |
| RORγT | AFKJS-9 | PE | Transcriptional marker | Intranuclear |
| PLZF1 | R17-809 | PE CF594 | Transcriptional marker | Intranuclear |
| Fox3P | 150D | AF488, AF647 | Transcriptional marker | Intranuclear |
| HLADR | G46-6 | BV650 | T cell activation | Surface |
| CD39 | eBioA1 (A1) | FITC, PE Cy7 | T cell activation | Surface |
| CD25 | BC96, M-A251 | PE, BV605 | T cell activation | Surface |
| CD69 | TP1.55.3 | ECD | T cell activation | Surface |
| ki67 | B56 | AF647 | T cell proliferation | Intracellular |
| TIGIT | MBSA43 | FITC, PerCP EFluor710 | T cell activation/exhaustion | Surface |
| PD1 | EH12.2H7 | BV605 | T cell activation/exhaustion | Surface |
| IFNγ | 4S.B3 | FITC, BV510 | Cytokine | Intracellular |
| TNFα | Mab11 | AF700, PerCP Cy5.5 | Cytokine | Intracellular |
| CD107a | H4A3 | APC, BV605 | Degranulation | Surface |
| LIVE/DEAD | ---- | Near IR | Dead cell stain | ---- |

### SIV Viral RNA quantification

Viral RNA was isolated from necropsy tissue homogenates by adding TriZol and extracting RNA via a standard phenol-chloroform extraction. Viral RNA was quantified using a *gag* qPCR assay as previously described [41]. Viable cell counts were determined from an aliquot of the same homogenate and used to calculate the SIV copies per cell. If the cell count was below the limit of detection of the hemocytometer, the limit of detection divided by two was used to approximate the value.

### Mtb barcode sequencing

Mtb genomic DNA was isolated as previously described [22]. Following gDNA isolation, samples were quantified and diluted to 10ng/uL. Samples were then amplified twice using 2x Q5 Master Mix (New England BioLabs) and two unique primer sets, one to add a molecular counter, and one to add the Illumina TruSeq adapter sequences. Primer sequences can be found in Supplemental Table 2. Samples were then sequenced on an Illumina MiSeq using a 2×150 kit and v2 chemistry. A computational pipeline courtesy of MRC and the Fortune lab was used to determine barcode sequences. Barcode diversity was then determined using Simpson’s Diversity index, and comparisons between groups were done using Kruskal-Wallis. All figures were generated using Prism 8 and Adobe Illustrator 2019.

### Statistical Analysis

All total CFU counts were transformed by adding 1 to account for sterile granulomas and LN tissues in log-scale graphs. Statistics were done using Prism 8. Comparisons between groups were done using either Mann-Whitney, for comparisons between two groups, or Kruskal-Wallis, for comparisons between three groups. Spearman correlation coefficients were calculated to determine relationships between variables.

## Acknowledgements

These studies were supported by NIH RO1 AI-111815 and NIH R21 AI127127. We thank the animal care staff at the University of Pittsburgh for excellent care of the animals housed at these facilities. Research reported in this publication was supported in part by the Office Of The Director, National Institutes of Health under Award Number P51OD011106 to the WNPRC, University of Wisconsin-Madison. The content is solely the responsibility of the authors and does not necessarily represent the official views of the National Institutes of Health. Finally, we would also like to thank Virology Services, a member of Research Services, at the WNPRC for performing SIV viral load assays.

---

**Figure S1.**
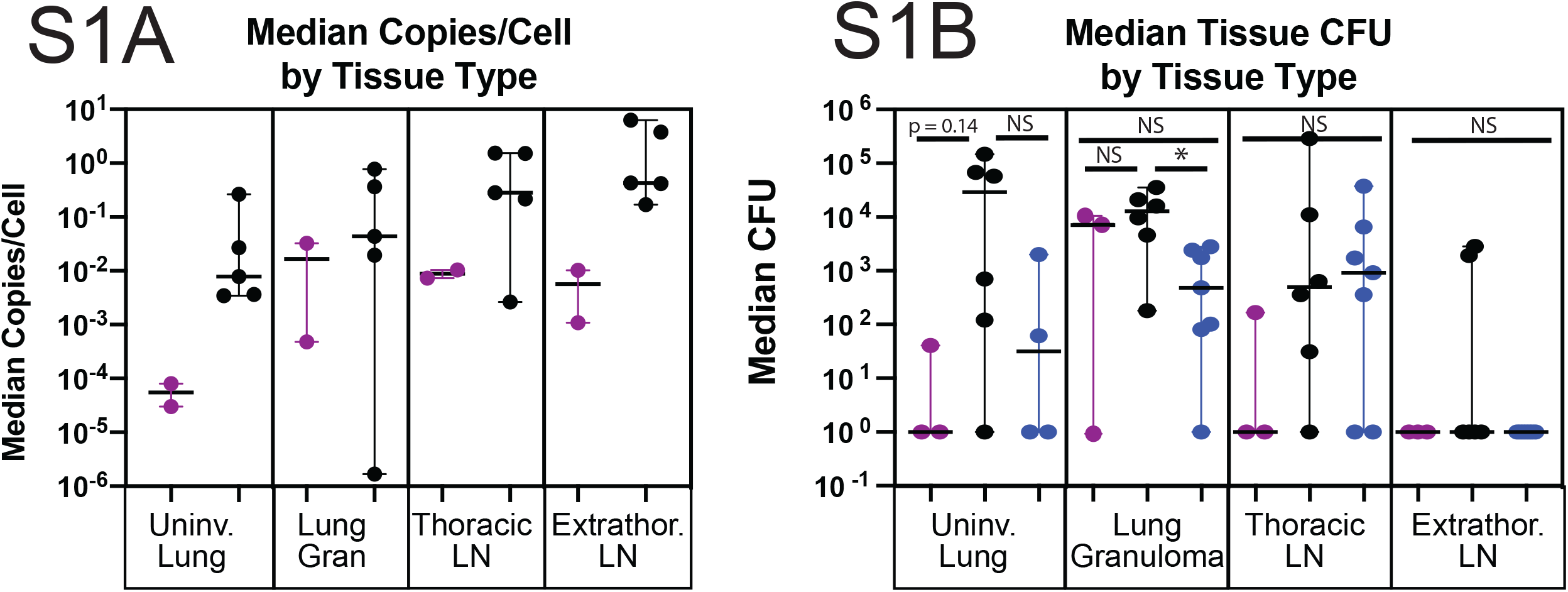
Animal medians of copies/cell (A) and bacterial CFU (B) by tissue type and controller status. Purple indicates animals able to spontaneously control SIV replication below 1000 SIV copies/mL of plasma, black indicates animals that could not control SIV replication, and blue indicates SIV-naïve animals. Significance determined by Kruskal-Wallis: *, p < 0.05, **, p <0.01, ***, p < 0.001, ****, p < 0.0001.

**Figure S2.**
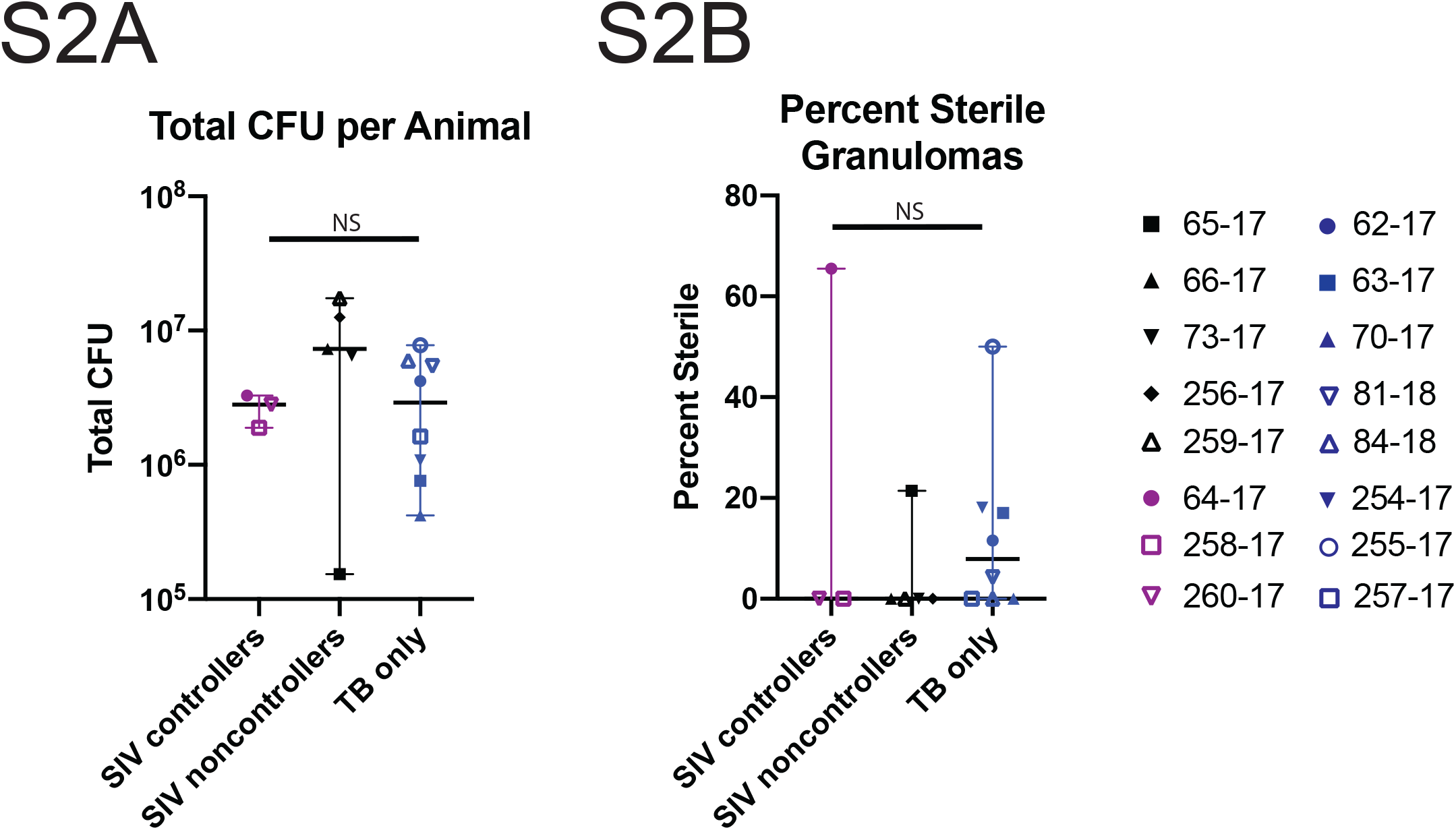
Total CFU per animal (A) and percent sterile granulomas (B) for each animal and cohort. Significance determined by Kruskal-Wallis.

**Figure S3.**
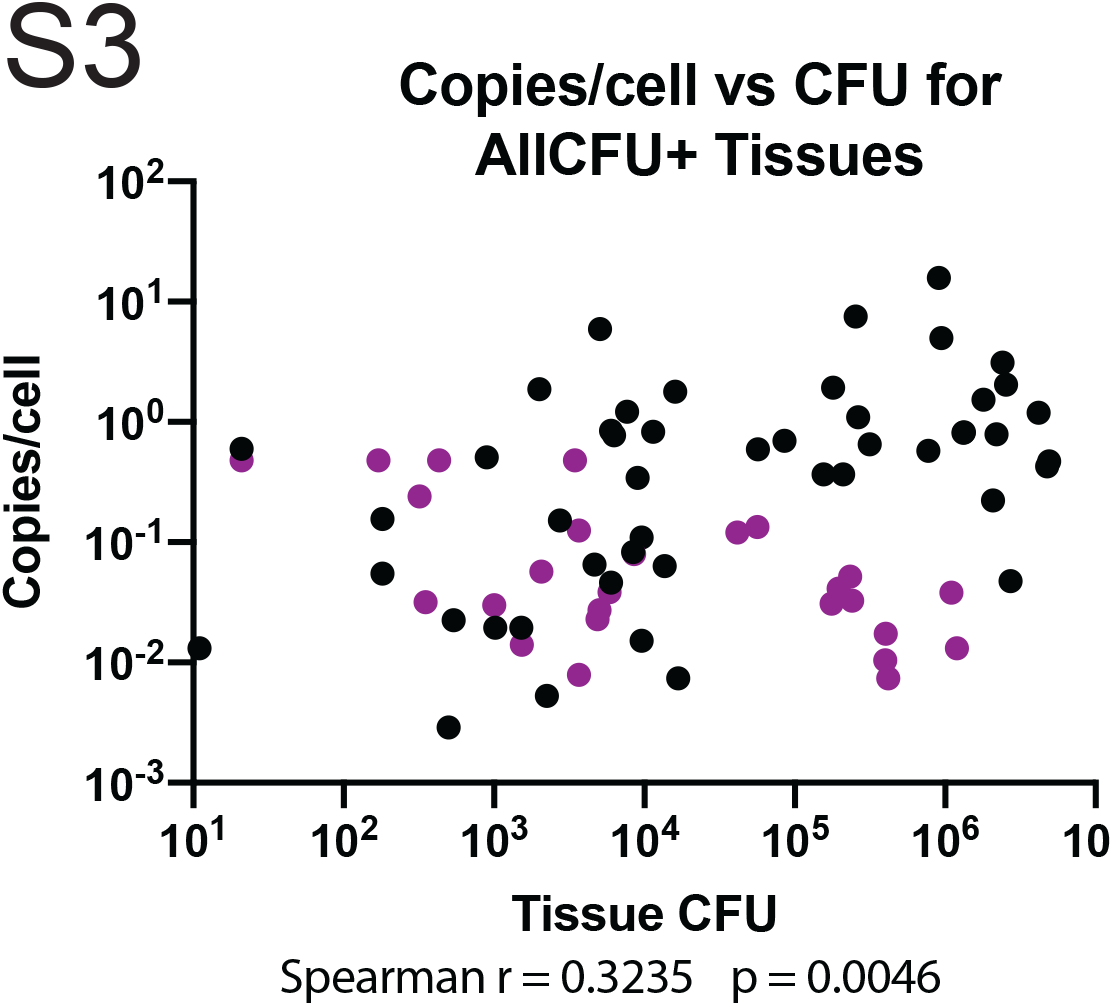
SIV present in individual lesions measured by copies/cell vs bacterial CFU per lesion in lung granulomas, thoracic LNs, and lung tissue for SIV non-controllers (black) and SIV controllers (purple). Spearman correlation coefficients were done to determine significance. Correlations were considered significant if p < 0.05.

**Figure S4.**
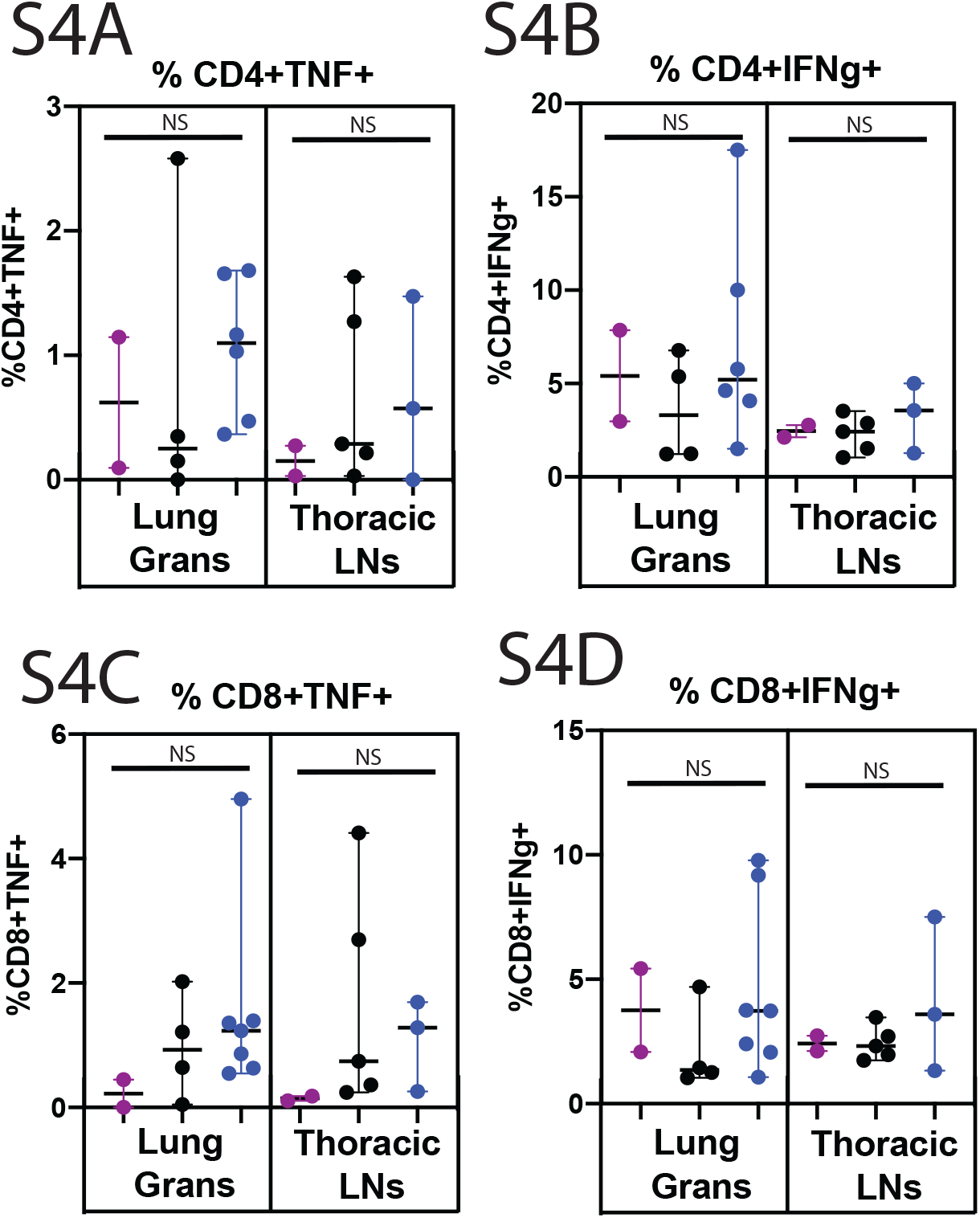
Animal medians of CD4+ (A, B) and CD8+ (C,D) T cells producing TNFα (A, C) or IFNγ (B, D) in the lung granulomas (left panel) or thoracic LN (right panel) in SIV controllers (purple), SIV non-controllers (black), and SIV-naïve (blue) animals. Each dot represents a single animal. Significance determined by Kruskal-Wallis: *, p <0.05, **, p < 0.01, ***, p <0.001, ****, p < 0.0001.

**Figure S5.**
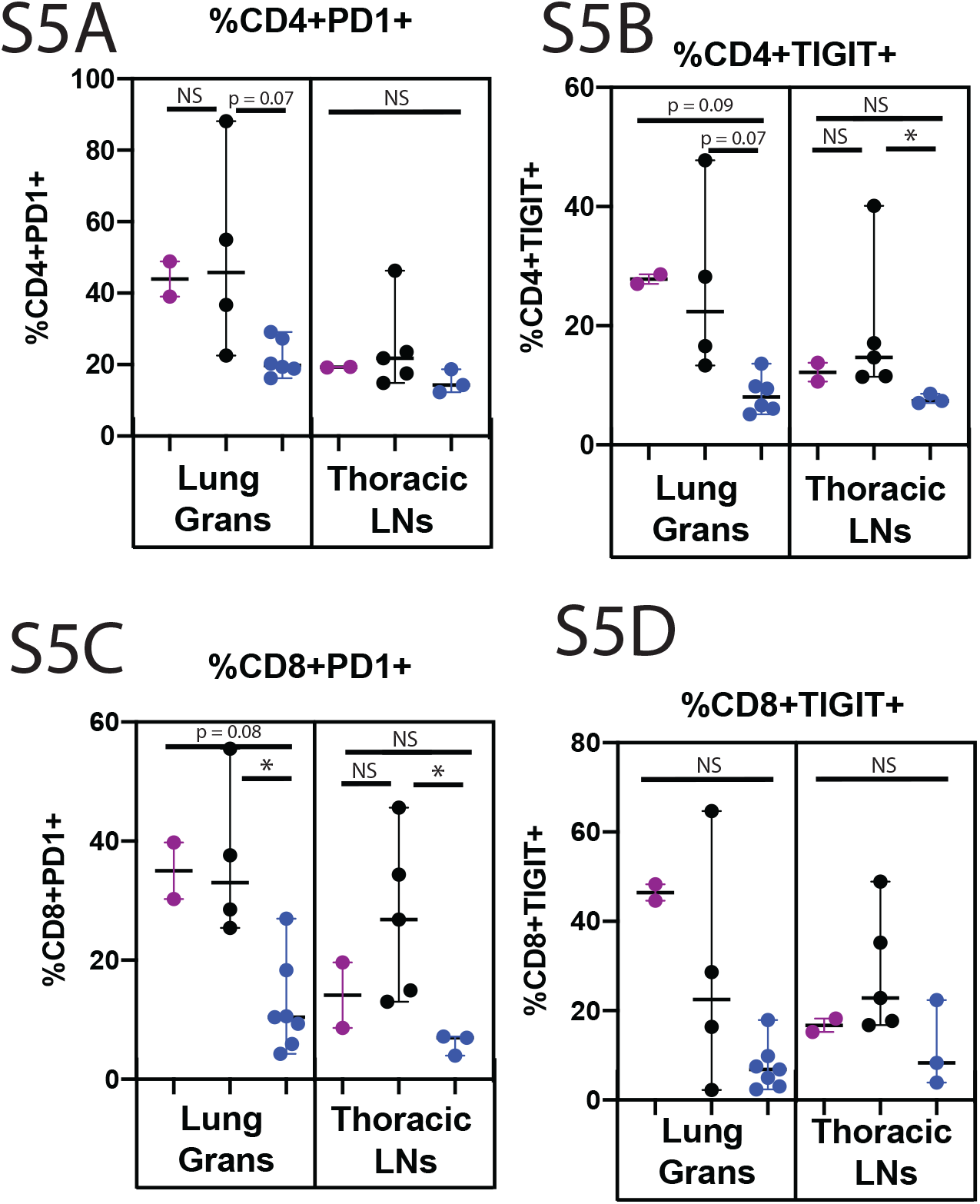
Animal medians of CD4+ (A, B) and CD8+ (C,D) T cells expressing activation markers PD1 (A, C) or TIGIT (B, D) in the lung granulomas (left panel) or thoracic LN (right panel) in SIV controllers (purple), SIV non-controllers (black), and SIV-naïve (blue) animals. Each dot represents a single animal. Significance determined by Kruskal-Wallis: *, p <0.05, **, p < 0.01, ***, p <0.001, ****, p < 0.0001.

**Figure S6.**
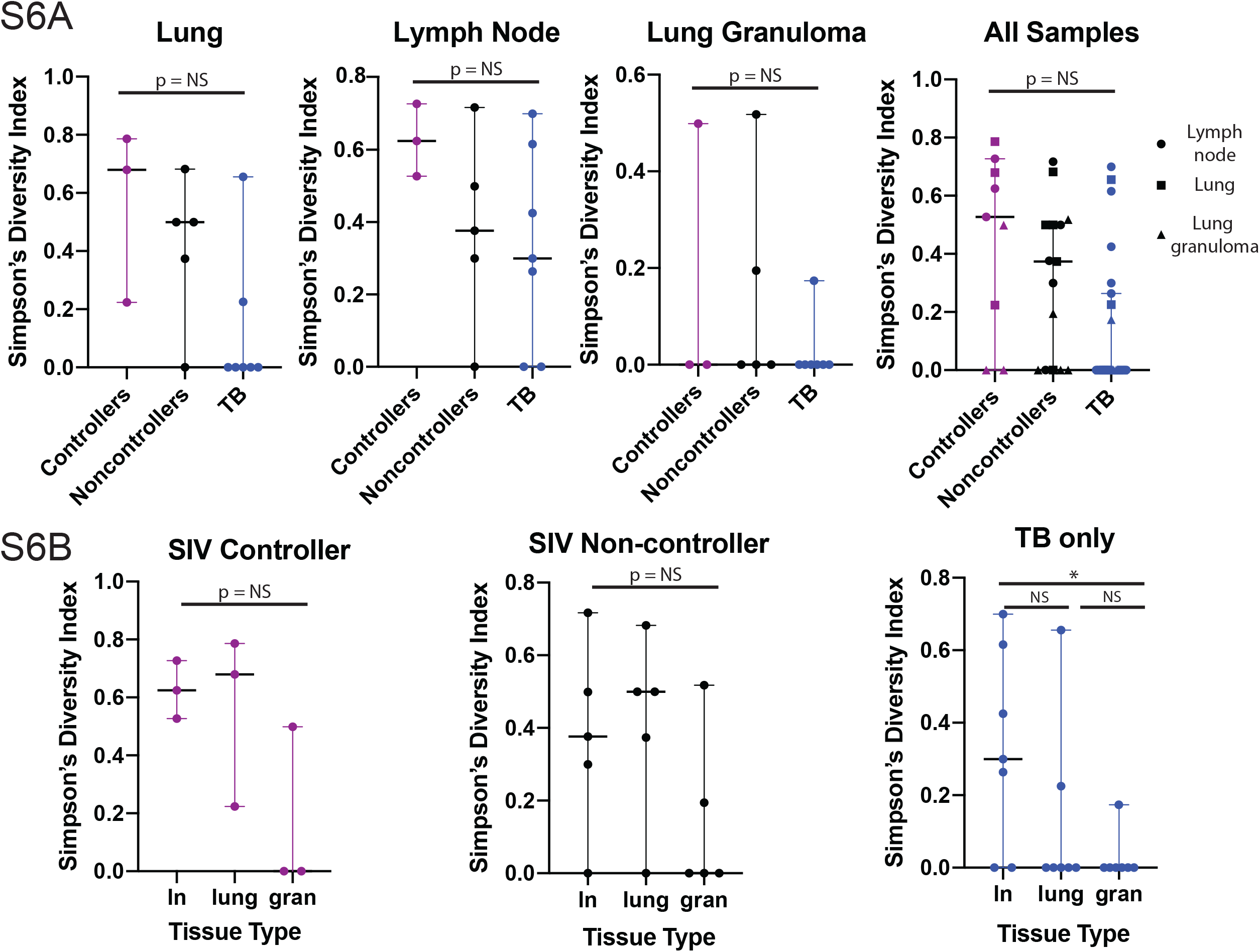
Animal medians of Mtb barcode diversity as calculated by Simpson’s Diversity Index in uninvolved lung, LN, and granulomas (A), as well as between SIV controllers (purple), SIV non-controllers (black), and SIV-naïve (blue) animals (B). Significance is determined by Kruskal-Wallis: *, p < 0.05, **, p < 0.01, ***, p < 0.001, ****, p < 0.0001.

**Supplemental Table 1.**
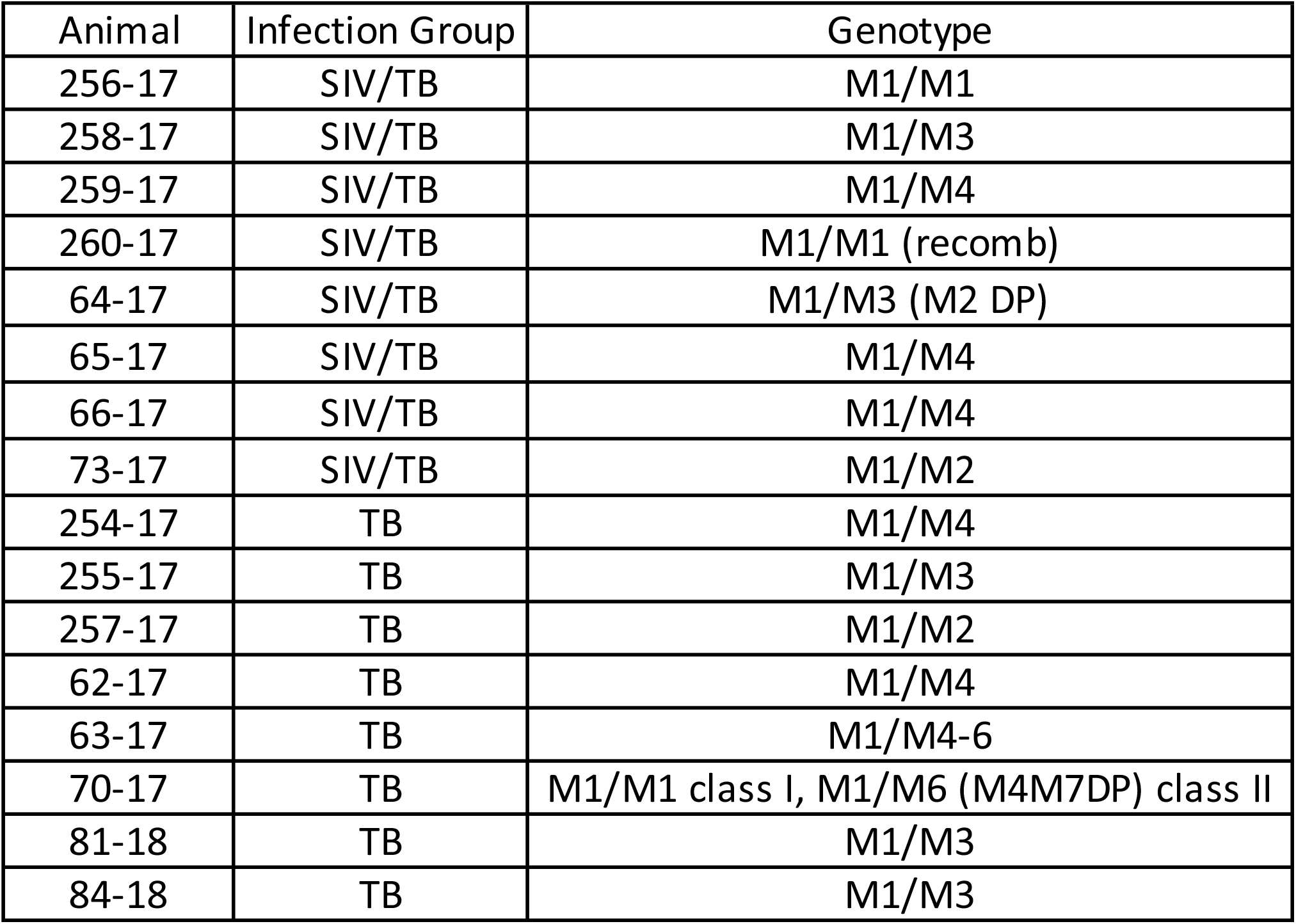

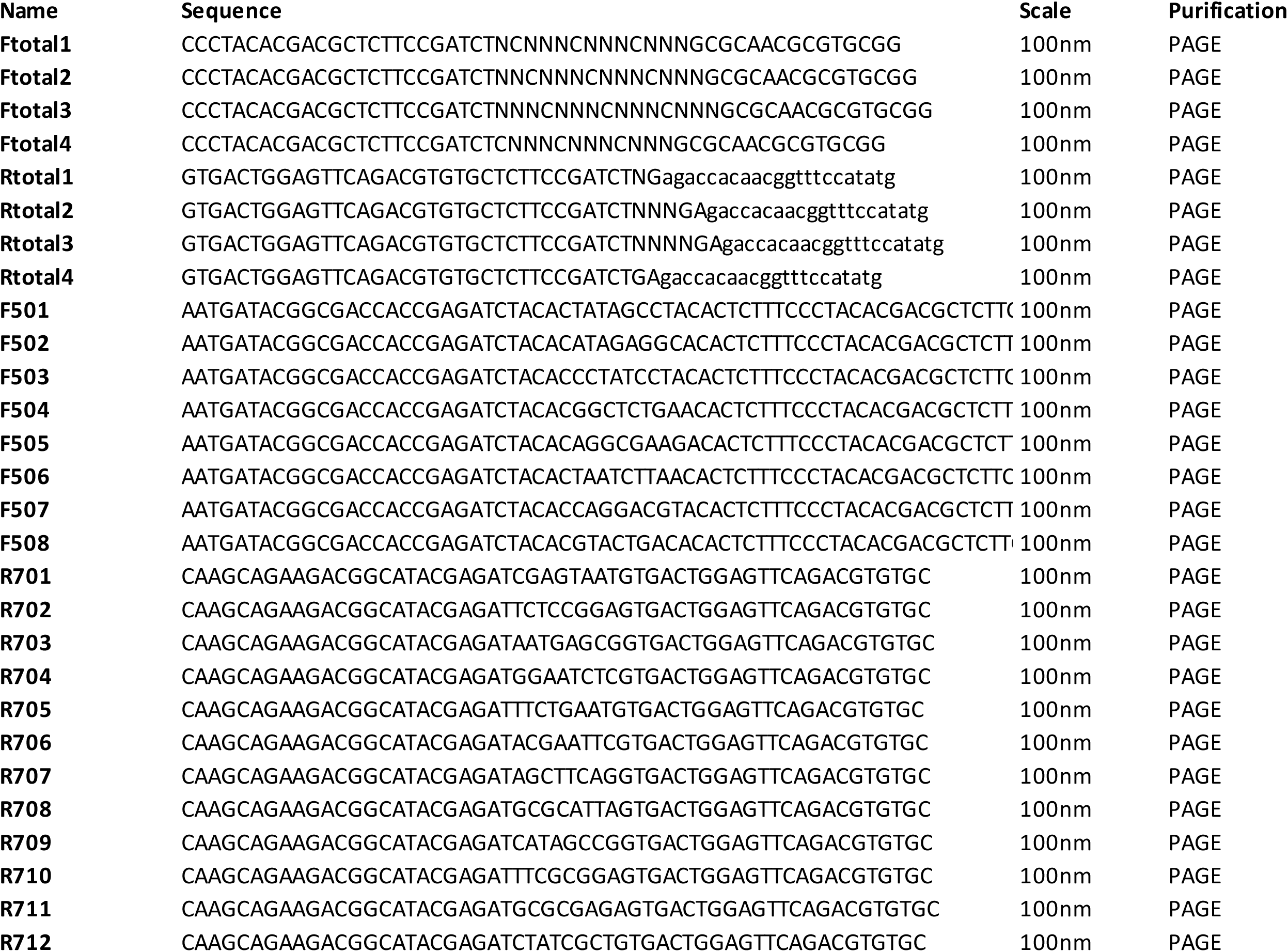

